# REMBRANDT: A high-throughput barcoded sequencing approach for COVID-19 screening

**DOI:** 10.1101/2020.05.16.099747

**Authors:** Dario Palmieri, Jalal Siddiqui, Anne Gardner, Richard Fishel, Wayne O. Miles

**Author notes:** Corresponding authors: DP, RF and WOM.

## Abstract

The Severe Acute Respiratory Syndrome Coronavirus-2 (SARS-CoV-2), also known as 2019 novel coronavirus (2019-nCoV), is a highly infectious RNA virus. A still-debated percentage of patients develop coronavirus disease 2019 (COVID-19) after infection, whose symptoms include fever, cough, shortness of breath and fatigue. Acute and life-threatening respiratory symptoms are experienced by 10-20% of symptomatic patients, particularly those with underlying medical conditions that includes diabetes, COPD and pregnancy. One of the main challenges in the containment of COVID-19 is the identification and isolation of asymptomatic/pre-symptomatic individuals. As communities re-open, large numbers of people will need to be tested and contact-tracing of positive patients will be required to prevent additional waves of infections and enable the continuous monitoring of the viral loads COVID-19 positive patients. A number of molecular assays are currently in clinical use to detect SARS-CoV-2. Many of them can accurately test hundreds or even thousands of patients every day. However, there are presently no testing platforms that enable more than 10,000 tests per day. Here, we describe the foundation for the REcombinase Mediated BaRcoding and AmplificatioN Diagnostic Tool (REMBRANDT), a high-throughput Next Generation Sequencing-based approach for the simultaneous screening of over 100,000 samples per day. The REMBRANDT protocol includes direct two-barcoded amplification of SARS-CoV-2 and control amplicons using an isothermal reaction, and the downstream library preparation for Illumina sequencing and bioinformatics analysis. This protocol represents a potentially powerful approach for community screening, a major bottleneck for testing samples from a large patient population for COVID-19.

## Introduction

COVID-19 is an infectious disease whose etiopathogenic agent is the Severely Acute Respiratory Syndrome Coronavirus 2 (SARS-CoV-2) RNA virus (1). To date, this viral infection has globally affected over 4.5 million of patients and claimed more than 300,000 lives (2). The enormous volume of COVID-19 patients has placed significant strain on healthcare systems around the world and led to the confinement of over half of the world’s population.

Different regions of the world are enduring different stages in the battle against COVID-19. These range from initial identification of acute COVID-19 cases to preparing to re-open schools and businesses. Regardless, rapid large-scale screening for COVID-19 is essential. At present, the efficacy of seroconversion tests is still to be determined. Moreover, seroconversion following a SARS-CoV-2 infection displays an as yet unknown protective immunity (3). Thus, detection of the viral RNA genome remains the best predictor of both early infection, pre- or asymptomatic stages as well as an indicator of viral clearance and reduced or eliminated viral transmissibility. To accommodate state- and country-wide populations that will require serial monitoring of 1-100 million residents, a scalable diagnostic test is required that may be established with widely available and existing equipment.

Here, we describe, and provide reduction-to-practice data of a mass COVID-19 screening platform: Recombinase Mediated BaRcoding/AmplificatioN Diagnostic Tool (REMBRANDT; Figure 1). This protocol contains a number of key advances that maximize output and sensitivity whilst retaining speed and efficiency. Specifically, REMBRANDT uses recombination and repair enzymes to detect and amplify the viral genomic RNA (4). The amplification simultaneously tags the individual samples with dual barcoded primers (Figure 2). This critical step rapidly generates DNA products that are significantly more stable than RNA. Utilizing two independent barcoded primers per well enables a single patient sample within a 96-well plate to be independently marked with a well-specific barcode *and* a plate-specific barcode. This algorithm reduces the number of barcodes 100-fold from conventional barcoding methods. Once barcoded, the patient samples from multiple 96-well plates can be pooled and purified, ultimately minimizing reagent usage, time and sample-to-sample variation. While the system is further scalable, we provide barcoded primers sufficient to distinguish individual samples for ninety six 96-well plates or 9,216 patient samples. Processing these samples together enables rapid library construction using any one of the barcoded Illumina kits. Utilizing 12 Illumina barcodes allows further sample multiplexing such that twelve different 9,216 combined samples may be further mixed together to perform Next Generation (NextGen) Sequencing analysis of 110,592 patient samples in a single run on a number of Illumina-based platforms. The smaller numbers of barcodes requires less time-consuming computational processing, as trimmed sequences can be quickly divided based on barcode, and mapped to SARS-CoV-2 N gene or a control human gene (RNAse P) sequences. Importantly, this protocol introduces a unique, synthetic SARS-CoV-2 N gene sequence into every 96-well plate. This control has identical primer annealing regions to the SARS-CoV-2 N gene but contains 6 engineered base pair substitutions that distinguishes it from native sequences; providing an internal quality control for the process and a measure of batch effects. The laboratory and computational framework is designed to maximize SARS-CoV-2 detection efficiency while minimizing reagent usage, processing and turn-around time.

**Figure 1:**
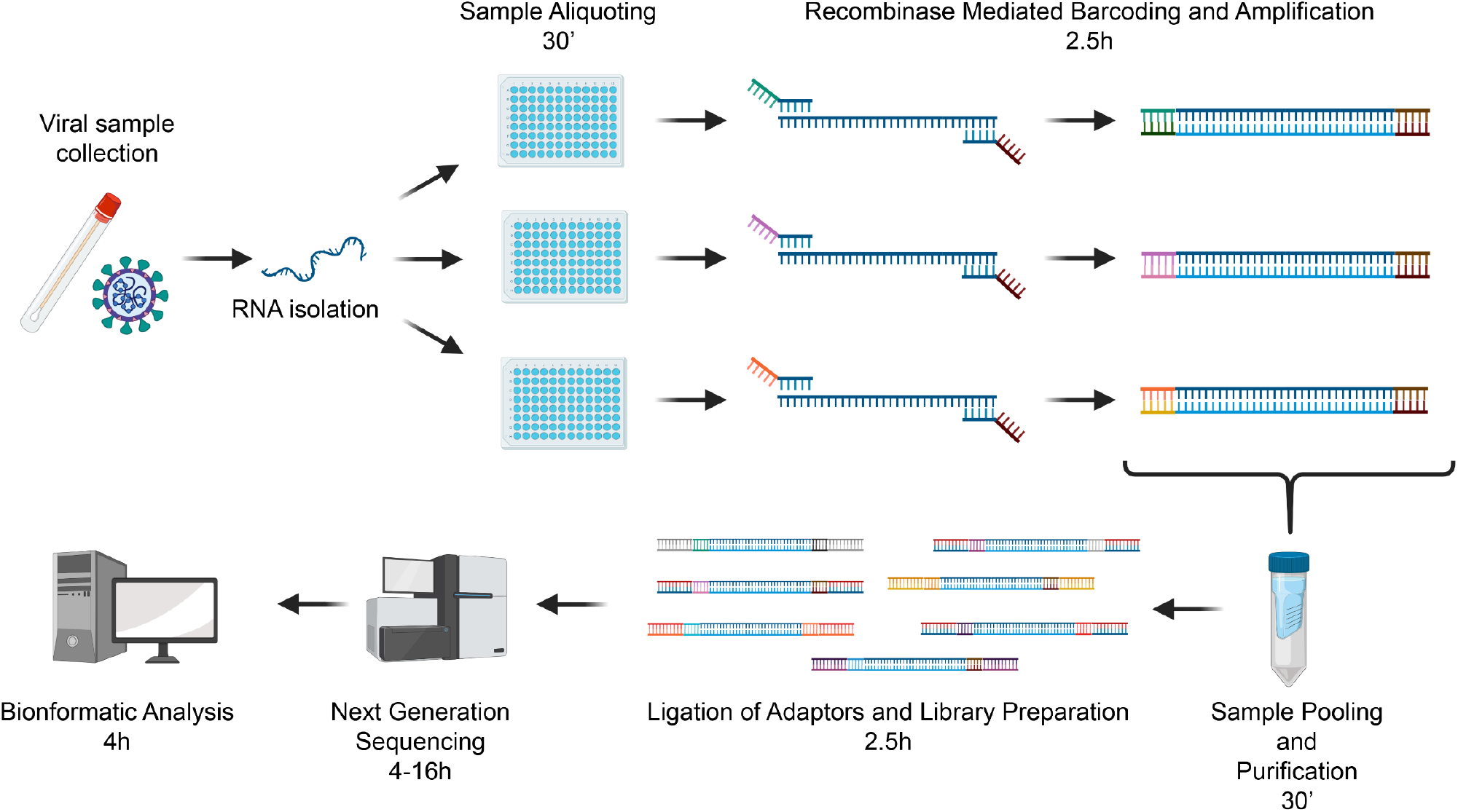
The REMBRANDT pipeline. Schematically detailing the protocol utilized for the REMBRANDT pipeline for testing for COVID-19.

**Figure 2:**
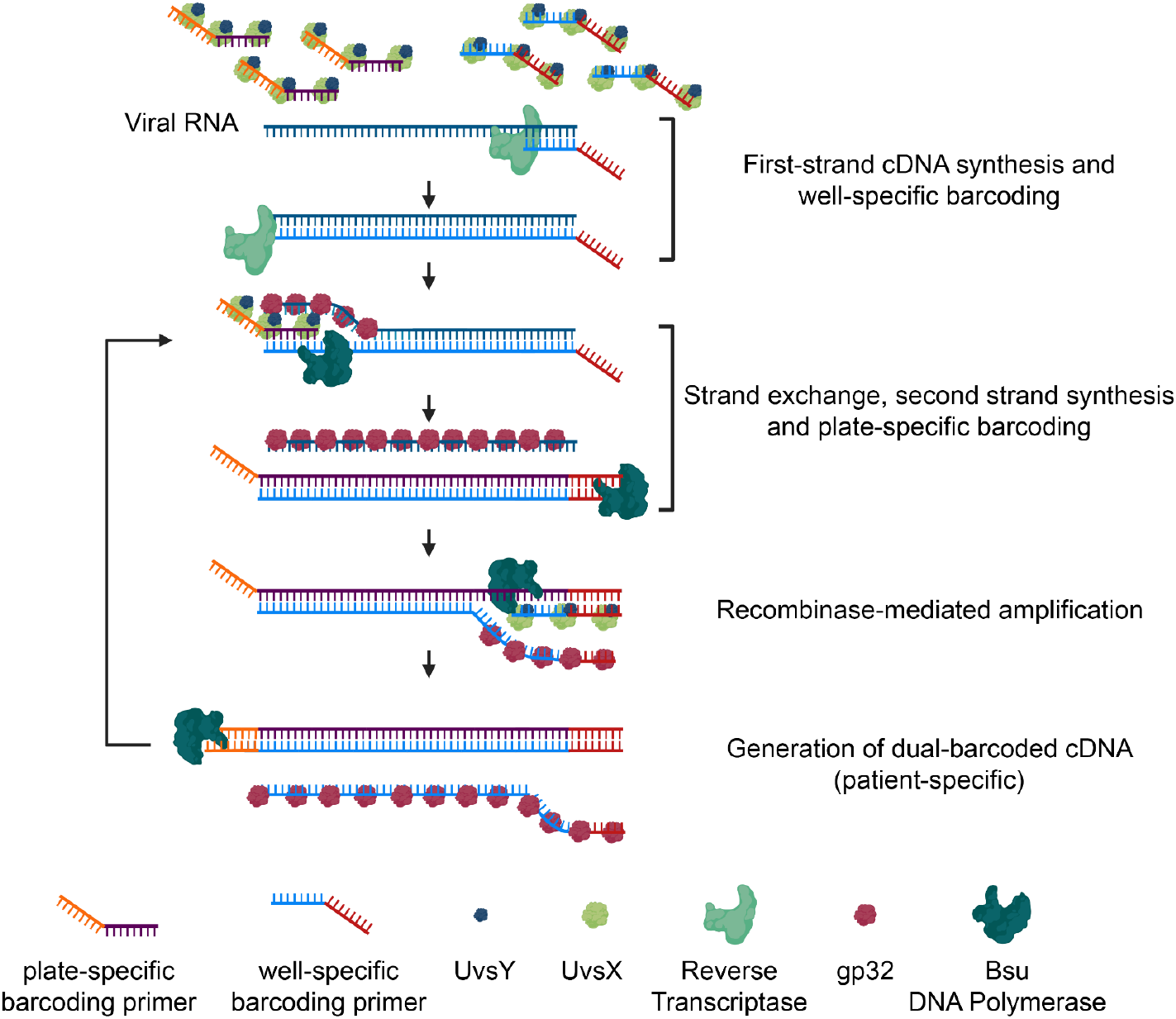
Isothermal Amplification and Barcoding strategy. Schematically detailing the protocol utilized for the isothermal amplification and dual-barcoding of target RNAs.

## Methods

### Starting Considerations

**1:** This protocol can effectively analyze 110,592 patient samples using 192 total barcoding primers (96 forward and 96 reverse) and 12 different adapters for High-Throughput sequencing. However, for ease of description the steps outlined here are defined for 9,216 samples or ninety six 96-well plates. To achieve this number, the protocol (steps RS1-8) should be performed 96 times. Each 96-well plate uses the same set of barcoded reverse primers to identify the patient and a separate barcoded forward primer to identify the plate. Each plate of 96 samples therefore has a different barcoded forward primer. This ultimately enables 12 different Illumina libraries to be combined and analyzed together for Next Generation Sequencing to reach the maximum potential of over 110,000 samples. To reach maximum throughput, the use of robotic aliquoting/pipetting system is recommended.

**2:** For this method, the RNA extraction from patients’ samples is not required. Based on previously reports, the protocol may be performed without anticipated issues using RNA extracted from nasopharyngeal swabs, following the CDC guidelines. However, recent manuscripts suggest that patients’ samples can be processed directly from nasopharyngeal swab, throat swab and saliva skipping the RNA extraction process (5). These direct detection approaches may reduce the number of amplification-available viral RNAs. Comparable results to commercial RNA extraction kits have been obtained using a 5-min direct detection preparation method of nasopharyngeal samples following 1:1 dilution with the Quick Extract DNA extraction Solution (Lucigen) (6). Moreover, different types of specimens, such as saliva, may be more sensitive for the detection of SARS-CoV-2 (7). We currently advise nasopharyngeal swabs as recommended by the US CDC. However, the saliva observations described above would appear to be the least invasive and easiest for patient collection.

**3:** RNA isothermal amplification can provide high sensitivity but are also prone to contamination. We recommend the use appropriate PPE, and the decontamination of benches to prevent RNA degradation. Sterile and barrier tips are recommended throughout, as is the separation of research spaces for the amplification, purification and library preparation steps.

**4:** Personnel safety must be a priority when performing any COVID-19 diagnostic approaches. Please refer to CDC guidelines for the most appropriate use of equipment and PPEs when working with SARS-CoV-2 and other respiratory viruses.

**Note:** A list of the reagents and their manufacturers used in our experiments is included in the Supplementary Information.

### Protocol

**In vitro RNA transcription:** to generate positive control RNAs, T7-flanked forward primers and standard reverse primers (Supplementary Information) were used in a PCR reaction to apply a region of each target gene with Q5 polymerase. Templates used in this work were COVID-19-N (10006625, IDT) and RNPase P (10006626, IDT). Gel purified PCR products (200ng) are purified and used in T7 (Roche 10881775001) *in vitro* transcription reaction. DNase is then added for 15mins, before the RNA is precipitated using standard ethanol RNA purification method (8).

### Plate Preparation (PP)

**PP1:** Chemically synthesize ninety six forward barcoding primers (plate-specific barcode) and ninety six reverse barcoding primers (well-specific barcode) (see sequences in Supplementary Information), dissolved at a 10 μM concentration in RNase-free water. A total of 2.8 ml of each primer is required for the full protocol (*includes 20% extra volume due to pipetting error*).

**PP2:** Aliquot 1 μl of every well-specific reverse primer into each well of a 96-well plate. Repeat this step for ninety six individual 96-well plates. This will generate ninety six plates that contain the same well-specific reverse barcoding primer in each respective well.

**PP3:** aliquot 1 μl of a single plate specific forward barcoding primer into each well of a single specific 96-well plate from PS2. Repeat this step with a unique forward plate specific barcoding primer per plate from PS2. *These preparation steps generate ninety six 96-well plates (9,216 unique wells), with each well containing one plate-specific forward primer and one well-specific reverse primer. PS2 and 3 can be repeated multiple times to optimize the procedure and aliquoted plates stored at −20C before use*.

### REMBRANDT Steps (RS)

**RS1:** Prepare Isothermal Amplification Buffer 2X (IAB2X).

**Table 1:** Isothermal Amplification Buffer 2X (IAB2X)

| Reagent | Stock concentration | Final Concentration | Required volume 96 samples<sup>#</sup> | Required volume 9,216 samples<sup>#</sup> |
| --- | --- | --- | --- | --- |
| Tris Acetate pH 7.8 | 1 M | 80 mM | 92.8 µl | 8.91 ml |
| K-acetate | 5 M | 200 mM | 46.4 µl | 4.45 ml |
| DTT | 1 M | 10 mM | 11.6 µl | 1.1 ml |
| ATP | 100 mM | 5 mM | 58 µl | 5.57 ml |
| dNTPs | 10 mM | 480 µM | 55.68 µl | 5.35 ml |
| PEG-3500 | 44% | 11 % | 290 µl | 27.84 ml |
| Trehalose | 40% | 10 % | 290 µl | 27.84 ml |
| Phosphocreatine | 0.5 M | 50 mM | 116 µl | 11.13 ml |
| Creatine Kinase | 2 µg/µl | 200 ng/µl | 116 µl | 11.13 ml |
| Acetylated BSA | 10mg/ml | 200 µg/ml | 23.2 µl | 2.23 ml |
| H2O | / | / | 60.32 µl | 5.79 ml |
| Final volume |  |  | 1.16 ml | 111.36 ml |

Keep the 2X IAB2X mix on ice.
*<sup>#</sup>Note: 10 µl are required for each sample. Values are referred to 96 or 9,216 samples + 20% extra due to pipetting error (116 and 11,136 samples, respectively)*

**RS2:** Prepare the UvsX/UvsY mix by combining 10 ml of recombinant UvsX (5 mg/ml) and 10 ml of recombinant UvsY (2mg/ml). Add 2 μl of the resulting mix to each well of the 96 well plates. We recommend gentle trituration 2-3 times of the UvsX/UvsY mix with the primers. This premixed plate should be kept on ice for all subsequent additions.

**RS3:** Add 1 μl of template RNA to each well of the 96-well plates.

**RS4:** prepare the Enzyme Mix.

**Table 2:** Enzyme Mix.

| Reagent | Stock concentration | Final Concentration | Required volume 96 samples# | Required volume 9,216 samples# |
| --- | --- | --- | --- | --- |
| T4 Gene 32 Protein | 10 mg/ml | 0.25 mg/ml | 58 µl | 5.57 ml |
| Bsu DNA Polymerase | 5000 U/ml | 0.25 U/µl | 116 µl | 11.14 ml |
| RevertAid Reverse Transcriptase | 200 U/µl | 10 U/µl | 116 µl | 11.14 ml |
| SUPERase RNase Inhibitor | 20 U/µl | 1 U/µl | 116 µl | 11.14 ml |
| IAB2X | 2X | 1X | 1.16 ml | 11.14 ml |
| Water<sup>##</sup> |  |  | 58 µl | 5.6 ml |
| Total volume |  |  | 1.624 ml | 155.91 ml |

Keep the mix on ice. Proceed immediately to step RS5. <sup>##</sup>*Optional Note: water can be substituted with an equal amount of 20 nM control spike-in DNA, if used in the downstream analysis*

**RS5:** Dispense 14 μl of Enzyme Mix into the all wells of the of the 96-well plates. Keep on ice.

**RS6:** Add 1 μl/well of 280 mM MgOAc to start the reaction. Ideally, MgOAc should be dispensed on the side of the well, or under the well cap, and mixed with the sample by vortexing and/or immediate centrifugation. This approach allows a simultaneous reaction start for all the samples.

**RS7:** Place all the 96-well plates at 38° C for 2h.

**RS8:** Collect the REMBRANDT products from all the 96-well plates into a single container (each 96-well plate will yield 1.92 ml).

*Note: next steps are referred to the handling of a single 96-well plate, but can be scaled-up as needed. The maximum yield/well is about 1μg of amplification product. Since suggested purification columns have a maximum capacity of 500 μg, up to 5 plates can be purified with one column. Using a 20-spots vacuum manifold, the products of ninety six 96-well plates can be purified simultaneously*.

**RS9:** Add 5 volumes (9.6 ml) of DNA Binding Buffer. Mix well and load onto a Zymo-Spin VI column. Place the column(s) on a vacuum manifold, turn on the vacuum source and let the sample clear from the column (*repeat this step up to 5 times*). Add 5 ml of DNA wash Buffer. Repeat the wash step. Leave the vacuum source on for additional 5 minutes to remove all the wash buffer. Transfer the column into a 50 ml conical tube. Add 2 ml of water to the column and wait 1 minute. Centrifuge at 3,000 x g for 3 minutes and collect the eluted barcoded DNA.

**RS10:** If performing multiple plates/columns, combine the eluted barcoded DNA and label the combination as “Batch 1”

**RS11:** Repeat steps **RS1-10**. Ideally the procedure can be repeated a total of 24 times. The subsequent 1-24 Batches are individually used for library preparation, and SHOULD NOT be combined until AFTER LIGATION (Library construction) with batch specific adapters.

### Library construction (LC)

**LC1:** Before library construction, a fraction of each purified pool of barcoded products should be examined using the Bioanalyzer to determine concentration and product size. The respective size of the COVID-19 and RNase P product are 124 and 104 bp, respectively.

**LC2:** Using the Illumina library construction kit (e7490), start at protocol 1.6 and conduct End repair on each pool of DNA fragments.

**LC3:** Proceed to Adaptor Ligation Step

**LC4:** Proceed to Purification of Ligation of Reaction step. This is critical to remove any free adaptor that will add noise to the library construction.

**LC5:** PCR Enrichment of Ligation Reaction. We recommend using 8 cycles to minimize over-amplification.

**LC6:** Purification of PCR reaction.

**LC7:** The overall library quality should be checked using a Bioanalyzer to confirm the size of the DNA fragments. Libraries that contain detectable levels of Primer-Primer annealing events should be repeated.

**LC8:** Determine the absolute DNA concentration of each library using Qubit.

**Sequencing (S):** The libraries are then loaded onto the Illumina sequencing platform and run as per manufacturer’s instructions. Once complete, the dataset is downloaded onto a suitable storage server for analysis.

### Bioinformatics (B)

**B1:** De-multiplexing reads

- Create CSV files listing of all 3 sets of barcodes (Illumina, plate, well)
- Create CSV file listing combinations of barcode matched to a patient sample
- <u>Multiplexing Iteration</u>
- Build connection using the Fastq streamer function for file
- Loop over all chunks of reads
- Extract a chunk of reads using yield function
- Loop over set of barcodes
- *grepl* search for barcode
- If barcode present, write file to ‘fastq’ file
- End loop over barcodes
- End loop over all reads after all chunks are extracted
- Perform Multiplexing Iteration on initial file using Illumina barcodes
- Perform Multiplexing Iteration on resulting fastq files from Illumina barcode de-multiplexing using plate-specific barcodes
- Perform Multiplexing interaction on resulting files from de-multiplexing plate-specific barcodes using well-specific barcodes
- Match files to patient samples and perform mapping via RSubread as below

**B2:** Build index for mapping reads using RSubread

- Create reference fasta file from nCoV, Human RPP30, and SARS-CoV-2 sequences
- Use buildIndex function to construct index
- ref being location of fasta file

buildindex(basename=“./reference_index”,reference=ref)

**B3:** Construct SAF annotation file for RSubread alignment and featureCounts formatted as below:

GeneID Chr Start End Strand

**B4:** Map de-multiplexed reads to reference index using align() function

> align.stat <-align(index = “./reference_index”, readfile1 = reads1, readfile2 = reads2, output_file = “./Rsubread_alignment.BAM”)

**B5:** Obtain counts using *featureCounts()*

- featureCounts function on resulting BAM files with SAF annotation file

## Results

To establish the REMBRANDT protocol, whilst minimizing virus exposure, we amplified a region of the SARS-CoV-2 N gene and the control human RNase P gene (RPP30) using PCR (Figure 3A) using commercially available templates (COVID-19-N: 10006625, IDT and RNPase P: 10006626, IDT). Each of the forward primers contained an extended region that coded for the T7 polymerase binding site. This site was then used to synthesize each RNA using T7 polymerase transcription that was then purified using ethanol precipitation. We then utilized different dilutions of the RNase P RNA to identify specific conditions for the accurate and reproducible amplification of this region using a combined Reverse Transcriptase-Recombinase-Polymerase Amplification (RT-RPA) (Figure 3B). The optimized RT-RPA conditions were then evaluated for the SARS-CoV-2 N amplicon, which were found to efficiently amplify this region (Figure 3C). These results demonstrate that the REMBRANDT approach can directly rapidly amplify RNA from the CDC-recommended control gene and SARS-CoV-2 N gene regions.

**Figure 3:**
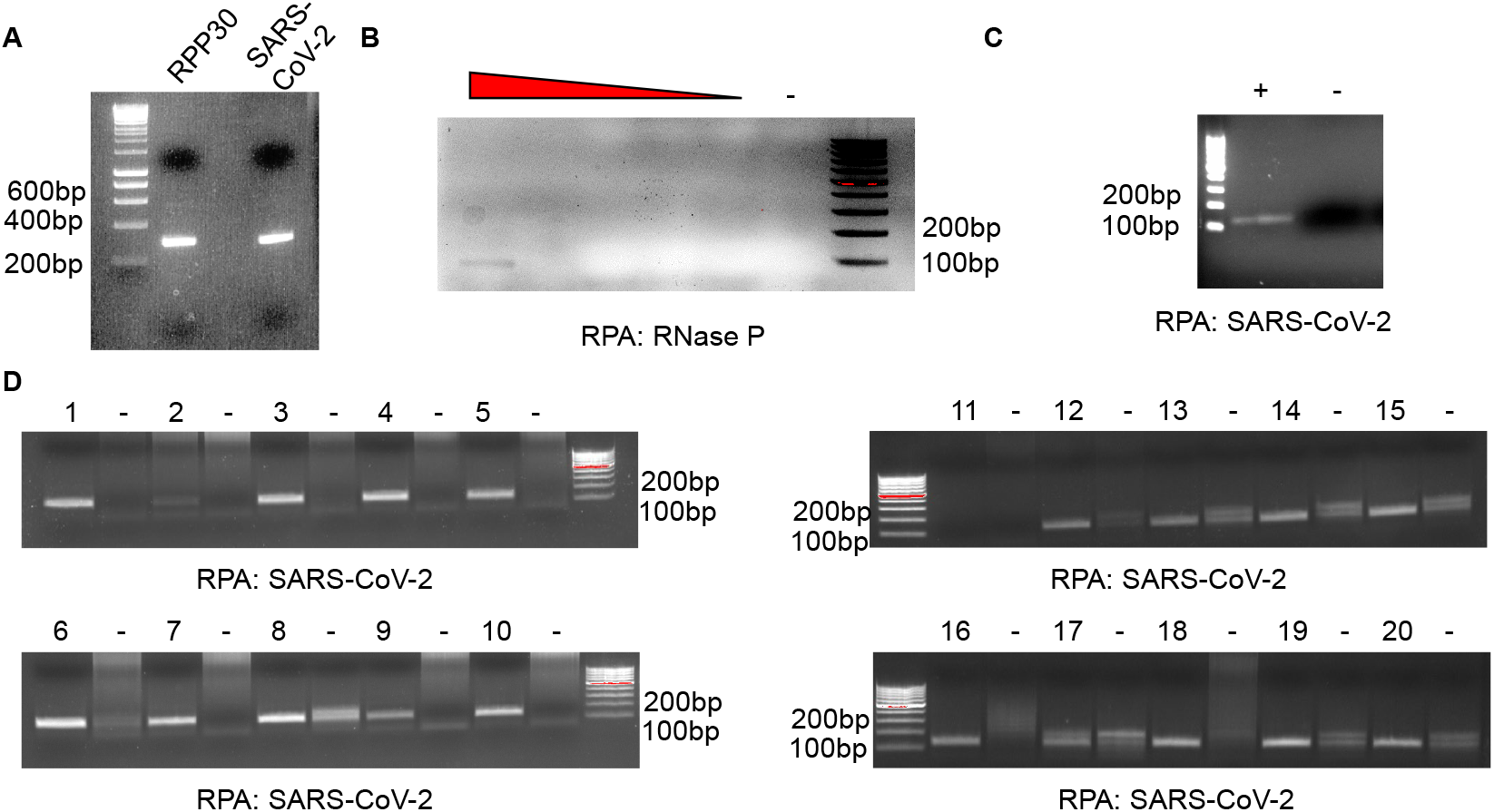
Isothermal Amplification of SARS-CoV-2 and RNase P RNAs. **(A)** Agarose gel of the T7 PCR products of a region of the SARS-CoV-2 N gene and human RNAse P. **(B)** Agarose gel of the isothermal amplification of human RNAse P from purified RNA or control (water, -). **(C)** Agarose gel of the isothermal amplification of SARS-CoV2-N gene from purified RNA or control (water, -). **(D)** Agarose gels using 8 different forward and 12 different reverse barcoded primers for the isothermal amplification of the SARS-CoV-2 N gene. Numbered lanes represent unique barcode combinations; – represents water control for each primer pair.

We then examined different combinations of barcoded SARS-CoV-2 N gene primers. We found that 8 unique barcoded forward and 12 unique barcoded reverse primers efficiently amplified the COVID-19 region. From this analysis, we determined that our test primer combinations could be utilized to detect the SARS-CoV-2 RNA (Figure 3D). Following the protocol described above, these barcoded DNA fragments can be rapidly assembled into a library for standard RNA-sequencing (RNA-Seq).

Unlike normal RNA-Seq runs that focus on mapping a small number of barcodes to a large number of genes, our computational strategy simply demultiplexes large numbers of barcodes and then maps them to three genes (Figure 4). In brief, we have developed a computational framework that separates samples based on 3 sets of barcodes using a nested iteration from the *ShortRead* Bioconductor package (9). The RNA-Seq *fastq* files are first imported into the R environment using the *FastqStreamer()* function and the sequence information extracted using the *yield()* function in *FastqStreamer*. For each read iteration, we use the *grepl()* string searching function to identify input barcodes. If input barcodes are identified, the reads are placed in a new *fastq* file using the *writeFastq()* function. The new *fastq* files are the input files for the next round of iterations of inputting, string-searching, and splitting files using the next round of barcodes. These process iterations start by de-multiplexing based on Illumina barcodes followed by plate-specific barcodes followed by well-specific barcodes. This iterative approach can efficiently map reads in to each unique barcode combination. Once complete, all of the reads are mapped to the inputted control RNase P, SARS-CoV-2 N gene and control SARS-CoV-2 oligo using the *Rsubread* Bioconductor package (10). The sequences of each of these amplicons are then combined into a *fasta* file, and an index built using the *buildIndex()* function. A Simplified Annotation Format (SAF) file is also generated for these sequences. Using the *align()* function, alongside our assembled indexes, we align our demultiplexed ‘fastq’ files to the control SARS-CoV-2 N, Human RPP30, and SARS-CoV-2 N gene sequences. The counts per sequence are then summarized from the resulting BAM alignment files using the *featureCounts()* function alongside our SAF annotation file. We use the *parLapply()* function from the *snow* R package to run large numbers of samples in parallel to maximize efficiency (11). The steps for this pipeline are detailed in Figure 4.

**Figure 4:**
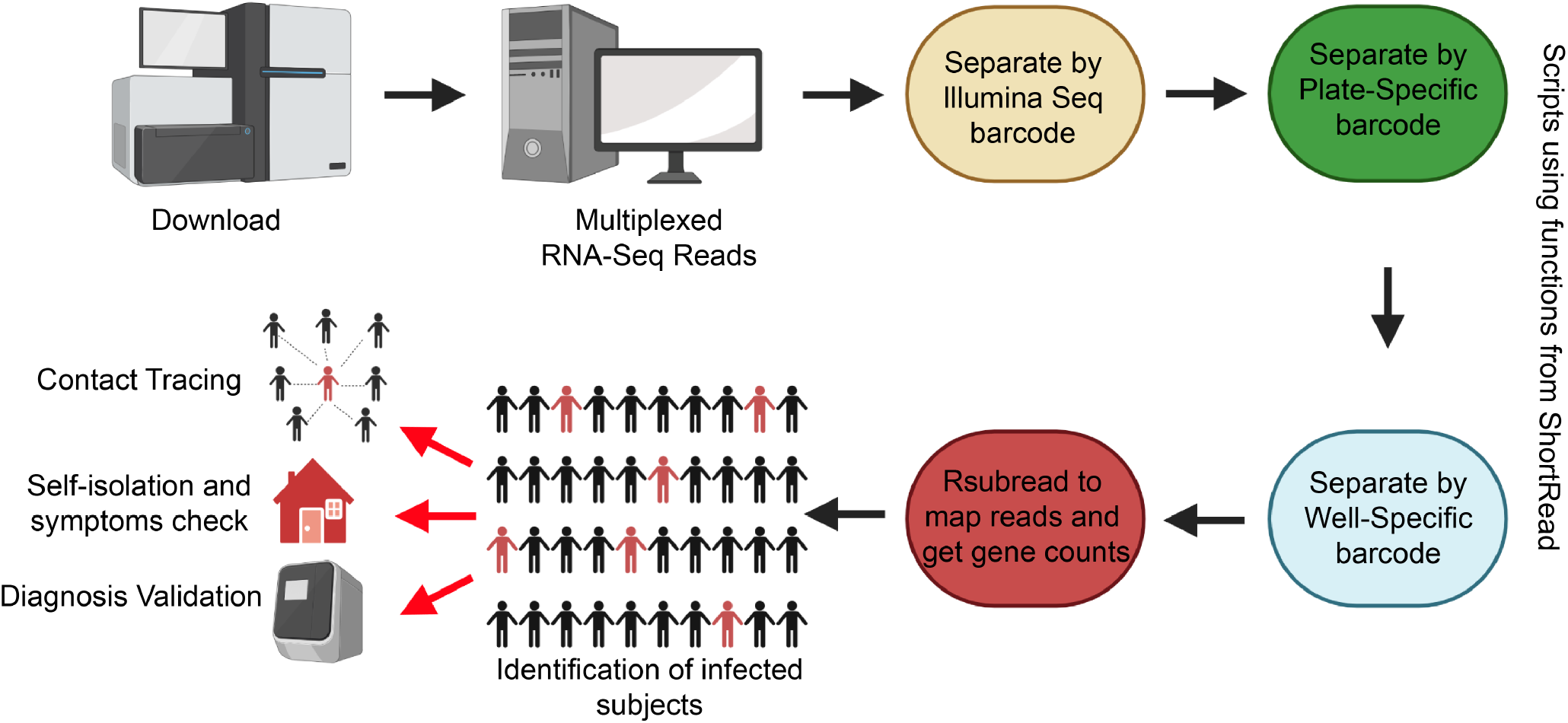
The computational framework for barcode and gene identification. Schematic detailing the protocol utilized for the bioinformatics analysis of the REMBRANDT RNA-seq data.

## Discussion and Conclusions

SARS-CoV-2 is a highly infectious single-stranded RNA virus. However, increasing evidence suggests that the vast majority of infected individuals display few or very mild symptoms (12). These people may still spread the virus for over 20 days after the initial contagion (13). For this reason, a key counter-measure against COVID-19 has been social distancing and city- or country-wide lockdowns. This approach has significantly slowed the number of new cases in the populations that practice these methods. Unfortunately, these lockdowns place real burdens on residents, and are designed to be temporary measures.

The rapid identification and quarantining of infected individuals is a key measure to containing the spread of SARS-CoV-2. Areas of the world such as South Korea and Northeastern Italy that have quickly tested their population *en masse* for SARS-CoV-2 have significantly reduced the spread of the disease. The relatively low incidence and mortality from these examples in comparison with adjacent countries and regions suggests that extensive population testing will be important for the foreseeable future, as cities and countries re-open their businesses and borders, to assess the COVID-19 status in real-time, at the community level (14).

One of the major current limitations of population testing is the availability of reagents (swabs, viral transport medium, RNA isolation kits, probes and other molecular biology reagents). The current clinical standard for COVID-19 diagnosis (qRT-PCR) requires suitable equipment for the amplification of the viral RNA and the detection of the infection. Each of these steps significantly slows sample processing and limits the number of tests that can be performed in a day. For this reason, new diagnostic tools are needed that are fast and efficient as well as scalable to population sized numbers and based on readily available reagents.

To address this need, the scientific community has delivered a remarkable and unprecedented number of assays to diagnose COVID-19. Several diagnostic approaches that can provide fast and cheap tests have been developed. One of the leading approaches uses blood tests to identify IgG and IgM antibodies against SARS-CoV-2 proteins (15). These tests show great promise. However, this assay can only examine the immunological status of the patient and cannot determine the viral load in exhaled respiratory particles, the major source of disease spread. These diagnostic kits will therefore be important to identify patients which may have developed immunity against the virus, but are not appropriate to test the ability of a patient to infect others.

A number of groups have been working to develop novel tools to detect the genetic material of SARS-CoV-2 within patient airways. These approaches allow the preliminary detection of potentially infected asymptomatic patients, whose diagnosis should be confirmed using standard qRT-PCR. The different tests range from the quick and portable kits (16–17), with the potential to become home-testing kits, to high-throughput multiplexed RT-PCR reactions. One particularly promising approach uses viral RNA reverse transcription and patient-specific barcoding of the single strand of cDNA, followed by cDNA amplification and NGS analysis (18). This approach, although interesting, requires 1 barcode per patient rather than multiplexing barcodes. Ultimately, that means to screen 10,000 patients one would require 10,000 barcodes. In addition, these large primers with large barcodes are likely to vary significantly in their amplification efficiency, making the individual testing of these primer sets essential.

REcombinase Mediated BaRcoding and AmplificatioN Diagnostic Tool (REMBRANDT) builds on these principles to double barcode each patient sample using an isothermal RT-RPA reaction. The combinatorial use of multiple forward and reverse barcodes, one per patient and one per plate, enables 192 primers to generate 9,216 patient-specific combinations. This number can then be further multiplexed and amplified with 12 Illumina barcodes utilized during library construction to examine 110,952 patient samples in a single Illumina run. This simplified and straightforward approach does not require specialized equipment for patient detection and library construction. We therefore have confidence that this system could be readily implemented in most communities, including those with limited resources, provided the availability of partners capable of NGS analysis. We are currently developing approaches to scale-up the production of the recombinant RT-RPA enzymes for the management of the potential increase in their demand.

As REMBRANDT uses an isothermal RNA reverse transcription and amplification reaction, it does not require PCR amplification. Moreover, since pairing of the template and primer during the amplification step relies on the activity of recombinases UvsX and UvsY, it is minimally affected by different *T_m_* or *T_a_* of the primers. The REMBRANDT pipeline also offers flexibility and can be readily adapted to detect other viral genes, viruses and/or other pathogenic species, by switching the design of the amplification primers. The protocol described here provides experimental evidence that the designed RT-RPA is effective in amplifying the synthetic SARS-CoV-2 N-gene RNA. Although the clinical efficacy of this approach remains untested, isothermal amplification has been previously used to detect SARS-CoV-2 levels in patients, from a number of bodily fluids (19).

The detailed protocol and experimental results for the fast (less than 24h) concurrent screening of over 100,000 potential COVID-19 samples as well as the framework to analyze NGS sequencing data is fully described. When clinically validated, REMBRANDT may represent a useful tool for screening easy-to-access bodily fluids for COVID-19 diagnosis.

## Acknowledgments

This work was supported by The Ohio State University Comprehensive Cancer Center (RF). The authors thank Dr. Kristine Yoder, Laura Miles and Anna Tessari for their critical support.

## Supplementary Data and code availability

Primer and oligonucleotides sequences used for this manuscript are detailed in Supplementary Information. Barcoded oligonucleotides to perform RT-RPA/barcoding on 100,000 simultaneous individual samples are also reported. REMBRANDT analysis code is deposited on Github (https://github.com/MilesLab/Rembrandt) and publicly available.

## References

1. Yuki K, Fujiogi M, Koutsogiannaki S. COVID-19 pathophysiology: A review. Clin Immunol. 2020 Apr 20;215:108427. doi: 10.1016/j.clim.2020.108427. PMID: 32325252 [Epub ahead of print]

2. https://www.worldometers.info/coronavirus/#countries (May 15th 2020)

3. Long QX, Liu BZ, Deng HJ, Wu GC, Deng K, Chen YK, Liao P, Qiu JF, Lin Y, Cai XF, Wang DQ, Hu Y, Ren JH, Tang N, Xu YY, Yu LH, Mo Z, Gong F, Zhang XL, Tian WG, Hu L, Zhang XX, Xiang JL, Du HX, Liu HW, Lang CH, Luo XH, Wu SB, Cui XP, Zhou Z, Zhu MM, Wang J, Xue CJ, Li XF, Wang L, Li ZJ, Wang K, Niu CC, Yang QJ, Tang XJ, Zhang Y, Liu XM, Li JJ, Zhang DC, Zhang F, Liu P, Yuan J, Li Q, Hu JL, Chen J, Huang AL. Antibody responses to SARS-CoV-2 in patients with COVID-19. Nat Med. 2020 Apr 29. doi: 10.1038/s41591-020-0897-1. [Epub ahead of print] PMID: 32350462

4. Piepenburg O, Williams CH, Stemple DL, Armes NA. DNA detection using recombination proteins. PLoS Biol. 2006 Jul;4(7):e204. PMID: 16756388

5. Emily A. Bruce EA, Huang ML, Perchetti GA, Tighe, Laaguiby P, Hoffman JJ, Gerrard DL, Nalla AK, Wei Y, Greninger AL, Diehl SA, Shirley DJ, Leonard DGB, Huston CD, Kirkpatrick BD, Dragon JA, Crothers JW, Jerome KR and Botten JW. Direct RT-qPCR detection of SARS-CoV-2 RNA from patient nasopharyngeal swabs without an RNA extraction step. doi: https://doi.org/10.1101/2020.03.20.001008 https://www.biorxiv.org/content/10.1101/2020.03.20.001008v2

6. Ladha A, Joung J, Abudayyeh OO, Gootenberg JS and Zhang F. A 5-min RNA preparation method for COVID-19 detection with RT-qPC. https://static1.squarespace.com/static/5b7c640be2ccd1703a3da4d3/t/5e8b5aec20703d6b23513142/1586191085082/Ladha+et+al.pdf

7. Wyllie AL, Fournier J, Casanovas-Massana A, Campbell M, Tokuyama M, Vijayakumar P, Geng B, Muenker MC, Moore AJ, Vogels CBF, Petrone ME, Ott IM, Lu P, Lu-Culligan A, Klein J, Venkataraman A, Earnest R, Simonov M, Datta R, Handoko R, Naushad M, Sewanan LR, Valdez J, White EB, Lapidus S, Kalinich CC, Jiang X, Kim DJ, Kudo E, Linehan M, Mao T, Moriyama M, Oh JE, Park A, Silva J, Song E, Takahashi T, Taura M, Weizman O, Wong P, Yang Y, Bermejo S, Odio C, Omer SB, Dela Cruz CS, Farhadian S, Martinello RA, Iwasaki A, Grubaugh ND and Ko AI. Saliva is more sensitive for SARS-CoV-2 detection in COVID-19 patients than nasopharyngeal swabs. doi: https://doi.org/10.1101/2020.04.16.20067835 https://www.medrxiv.org/content/10.1101/2020.04.16.20067835v1

8. Galgano A, Forrer M, Jaskiewicz L, Kanitz A, Zavolan M, Gerber AP. Comparative analysis of mRNA targets for human PUF-family proteins suggests extensive interaction with the miRNA regulatory system. PLoS One. 2008 Sep 8;3(9):e3164. doi: 10.1371/journal.pone.0003164. PMID: 18776931

9. Morgan M, Anders S, Lawrence M, Aboyoun P, Pagès H, Gentleman R. ShortRead: a bioconductor package for input, quality assessment and exploration of high-throughput sequence data. Bioinformatics. 2009 Oct 1;25(19):2607–8. doi: 10.1093/bioinformatics/btp450. Epub 2009 Aug 3. PMID: 19654119

10. Liao Y, Smyth GK, Shi W. The R package Rsubread is easier, faster, cheaper and better for alignment and quantification of RNA sequencing reads. Nucleic Acids Res. 2019 May 7;47(8):e47. doi: 10.1093/nar/gkz114.

11. Tierney L, Rossini AJ, and Li N. “Snow: A parallel computing framework for the R system.” International Journal of Parallel Programming 37.1 (2009): 78–90.

12. Li R, Pei S, Chen B, Song Y, Zhang T, Yang W, Shaman J. Substantial undocumented infection facilitates the rapid dissemination of novel coronavirus (SARS-CoV-2). Science. 2020 May 1;368(6490):489–493. doi: 10.1126/science.abb3221. Epub 2020 Mar 16. PMID: 32179701

13. He X, Lau EHY, Wu P, Deng X, Wang J, Hao X, Lau YC, Wong JY, Guan Y, Tan X, Mo X, Chen Y, Liao B, Chen W, Hu F, Zhang Q, Zhong M, Wu Y, Zhao L, Zhang F, Cowling BJ, Li F, Leung GM. Temporal dynamics in viral shedding and transmissibility of COVID-19. Nat Med. 2020 May;26(5):672–675. doi: 10.1038/s41591-020-0869-5. Epub 2020 Apr 15. PMID: 32296168

14. Angulo FJ, Finelli L, Swerdlow DL. Reopening Society and the Need for Real-Time Assessment of COVID-19 at the Community Level. JAMA. 2020 May 15. doi: 10.1001/jama.2020.7872. [Epub ahead of print] PMID:32412582

15. Venter M, Richter K. Towards effective diagnostic assays for COVID-19: a review. J Clin Pathol. 2020 May 13. pii: jclinpath-2020-206685. doi: 10.1136/jclinpath-2020-206685. [Epub ahead of print]. PMID 32404473

16. Broughton JP, Deng X, Yu G, Fasching CL, Servellita V, Singh J, Miao X, Streithorst JA, Granados A, Sotomayor-Gonzalez A, Zorn K, Gopez A, Hsu E, Gu W, Miller S, Pan CY, Guevara H, Wadford DA, Chen JS, Chiu CY. CRISPR-Cas12-based detection of SARS-CoV-2. Nat Biotechnol. 2020 Apr 16. doi: 10.1038/s41587-020-0513-4. [Epub ahead of print] PMID: 32300245

17. Joung J, Ladha A, Saito M, Segel M, Bruneau R, Huang MW, Kim N, Yu X, Li J, Walker BD, Greninger AL, Jerome KR, Gootenberg JS, Abudayyeh OO, Zhang F. Point-of-care testing for COVID-19 using SHERLOCK diagnostics. https://static1.squarespace.com/static/5b7c640be2ccd1703a3da4d3/t/5eb16fc2089aa30974484ce5/1588686798127/COVID-19+detection+%28v.20200505%29

18. Hossain A, Reis AC, Rahman S, and Salis HM. A Massively Parallel COVID-19 Diagnostic Assay for Simultaneous Testing of 19200 Patient Samples. https://docs.google.com/document/d/1kP2w_uTMSep2UxTCOnUhh1TMCjWvHEY0sUUpkJHPYV4/edit

19. Lamb LE, Barolone AN, Ward E, Chancellor MB. Rapid Detection of Novel Coronavirus (COVID-19) by Reverse Transcription-Loop-Mediated Isothermal Amplification. doi: https://doi.org/10.1101/2020.02.19.20025155. https://www.medrxiv.org/content/10.1101/2020.02.19.20025155v1

